# Semiparametric Confidence Sets for Arbitrary Effect Sizes in Longitudinal Neuroimaging

**DOI:** 10.1101/2025.02.10.637497

**Authors:** Xinyu Zhang, Kenneth Liao, Jakob Seidlitz, Maureen McHugo, Suzanne N. Avery, Anna Huang, Aaron Alexander-Bloch, Neil Woodward, Stephan Heckers, Simon Vandekar

## Abstract

The majority of neuroimaging inference focuses on hypothesis testing rather than effect estimation. With concerns about replicability, there is growing interest in reporting standardized effect sizes from neuroimaging group-level analyses. Confidence sets are a recently developed approach to perform inference for effect sizes in neuroimaging but are restricted to univariate effect sizes and cross-sectional data. Thus, existing methods exclude increasingly common multigroup or nonlinear longitudinal associations of biological brain measurements with inter- and intra-individual variations in diagnosis, development, or symptoms. We broadly generalize the confidence set approach by developing a method for arbitrary effect sizes in longitudinal studies. Our method involves robust estimation of the effect size image and spatial and temporal covariance function based on generalized estimating equations. We obtain more efficient effect size estimates by concurrently estimating the exchangeable working covariance and using a nonparametric bootstrap to determine the joint distribution of effect size across voxels used to construct confidence sets. These confidence sets identify regions of the image where the lower or upper simultaneous confidence interval is above or below a given threshold with high probability. We evaluate the coverage and simultaneous confidence interval width of the proposed procedures using realistic simulations and perform longitudinal analyses of aging and diagnostic differences of cortical thickness in Alzheimer’s disease and diagnostic differences of resting-state hippocampal activity in psychosis. This comprehensive approach along with the visualization functions integrated into the pbj R package offers a robust tool for analyzing repeated neuroimaging measurements.

## 1 Introduction

Brain-behavior associations use measurements obtained from magnetic resonance imaging (MRI) to study associations with diagnosis, symptomology, cognition, or other non-brain phenotypes. For example, brain-behavior associations may examine how MRI-derived hippocampal function is associated with diagnosis in early psychosis-spectrum disorders. Spatial extent inference is the most commonly used approach for investigating brain-behavior associations. This approach identifies regions of the brain image that are significantly associated with the non-brain phenotype by thresholding the statistical image for the voxel-level association above a given value, and, for each spatially contiguous “cluster”, computing a *p*-value representing the probability of observing a cluster of that size or larger if the phenotype was not associated with the brain measurement (Worsley et al., 1999).

In response to long-held concerns about replicability, there has been growing interest in reporting standardized effect size estimates to characterize the strength of findings in neuroimaging (Bowring et al., 2019, 2021; Kang et al., 2024; Marek et al., 2022; Owens et al., 2021; S. N. Vandekar & Stephens, 2021). Standardized effect sizes are unitless indices quantifying the strength of an association (Kelley & Preacher, 2012). However, spatial extent inference is strictly focused on hypothesis testing because it is defined under the null that the brain image is independent of the phenotype. Therefore, it is not amenable to performing inference for effect sizes, as they are only non-zero when the null is false. In contrast to spatial extent inference, performing effect size-based statistical inference has several advantages (Bowring et al., 2021; Sommerfeld et al., 2018; S. N. Vandekar & Stephens, 2021). First, it provides a more nuanced understanding of the magnitude and practical significance of findings, beyond the arbitrary classification using hypothesis tests. Second, it facilitates meta-analyses and comparisons across studies, which is particularly important in evaluating and improving replicability (Zimmermann et al., 2005).

Recent advances in methods for spatial “confidence sets” address some of the limitations of hypothesis testing-based approaches by constructing these sets for unstandardized effect sizes (Bowring et al., 2019) or specific effect sizes like Cohen’s *d* (Bowring et al., 2021). Confidence sets use a given effect size threshold to confidently identify two spatial regions that capture the region where the true effect size is above the threshold with a prespecified probability (Sommerfeld et al., 2018). Confidence set methods allow investigators, not only to reject a null hypothesis but to confidently identify regions of the brain where the effect size is very likely to be below a meaningful threshold (Ren et al., 2023). Existing confidence set methods are not applicable to multi-group comparisons or nonlinear associations with age or symptom severity, which are often used for modeling the complex associations in brain-behavior associations (Grabell et al., 2018; Zhou et al., 2023). In addition, they are not applicable to longitudinal data, which is increasingly valued for its ability to study within-subject associations (Kang et al., 2024; Van Os et al., 1999).

Here, we introduce a novel statistical framework that addresses the limitations of existing confidence set approaches using Generalized Estimating Equations (GEE) and the robust effect size index (RESI) (Guillaume et al., 2014; Jones et al., 2023; Liang & Zeger, 1986; S. Vandekar et al., 2020). Our approach allows for the construction of confidence sets for arbitrary effect sizes, including those relevant to the multi-dimensional parameters needed to study complex associations in brain-related disorders, while accounting for dependence between repeated measurements. We demonstrate the utility of our method through simulations, showing its robust performance in small sample sizes (≤ 200), and apply it to study effect sizes in longitudinal analyses of regional cortical thickness differences in Alzheimer’s disease and functioning in psychosis-spectrum disorders. Our results highlight the potential of this approach to provide new insights that are not possible with existing methods. The methods presented in this paper are built upon functions provided by the pbj R package (https://github.com/statimagcoll/pbj), and the codes for analyses can be found here (https://github.com/statimagcoll/Confidence_Sets.git).

## 2 Statistical Theory

### 2.1 Notation and background

Our confidence set methods are based on spatial extent inference in a longitudinal context (Friston et al., 1994; Guillaume et al., 2014). We let 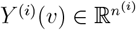 denote the outcome vector image of *n*^(*i*)^ repeated observations of brain measurements for *i* = 1, 2, ⋯, *n* independent subjects. These images are indexed by the voxel location, *v* ∈ 𝕍 ⊂ ℝ^3^, where 𝕍 denotes the bounded space of the brain, and we assume the data are registered to a template space. For each location *v*, we assume the model

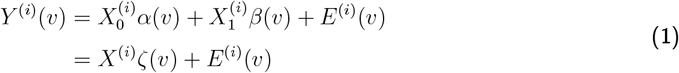

where 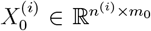 is a matrix of nuisance covariates including the intercept, 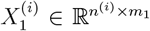 is a matrix of variables of interest (such as multi-group diagnosis or age fit with splines), *m* = *m*_0_ + *m*_1_, and 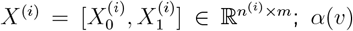 and *β*(*v*) are parameter image vectors that take values in 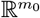 and 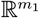, respectively, and 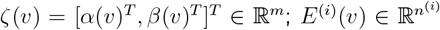 is an error term vector with mean zero and spatial covariance matrix Σ^(*i*)^(*v, w*) = Cov{*E*^(*i*)^(*v*), *E*^(*i*)^(*w*)} *<* ∞ for any two voxels *v* and *w*. This covariance describes the dependence between repeated imaging measurements over time and spatially within the image. The multilevel and spatial aspects of the data make them susceptible to unknown multivariate heteroskedasticity: Σ^(*i*)^(*v, w*) ≠ Σ^(*j*)^(*v, w*) for *i* ≠ *j*. To allow matrix notation, let *N* = Σ_*i*_ *n*^(*i*)^ be the number of all observations across *n* independent participants, 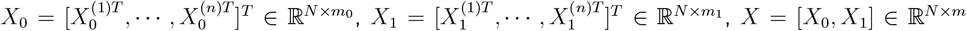, and *Y* (*v*) = [*Y* ^(1)*T*^ (*v*), ⋯, *Y* ^(*n*)*T*^ (*v*)]^*T*^ ∈ ℝ^*N*^ .

For estimation using GEE, let *V*_*w*_(*γ*) ∈ ℝ^*N×N*^ denote the block diagonal temporal working covariance matrix for all subjects, which we assume does not depend on the location in the image. For measurements from the same participant, *γ* controls the temporal covariance, e.g. exchangeable or AR1. Let *R*_0_ and *R* denote the symmetric *N × N* residual forming matrices for *X*_0_ and *X*, respectively (Supplementary Equation (S1)). To ease notations below, we do not explicitly include the dependence of *V*_*w*_ on *γ*.

### 2.2 Parameter estimation and test statistic

It is critical to account for repeated observations from the same participant when estimating *β*(*v*) and constructing confidence intervals. Prior methods for neuroimaging data only considered estimation using an independence working covariance structure due to the high computational demand of fitting over 100,000 voxel-level models (Guillaume et al., 2014). We perform estimation using working independence and exchangeable covariance structures using a one-step least squares estimator (Lipsitz et al., 2017). The one-step estimation speeds computation for the covariance parameters, which typically require iterative estimation in GEEs. The test statistic image is derived using robust covariance estimation in the context of GEEs treating each voxel separately (Huber, 1964; Liang & Zeger, 1986; White, 1980). Dependence among voxels is incorporated in preprocessing (via spatial smoothing) and in the effect size analysis using a joint bootstrap.

Based on model (1), we use the GEE

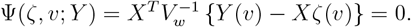

This estimating equation yields the unbiased least squares estimator for *β*(*v*) (S. N. Vandekar et al., 2019),

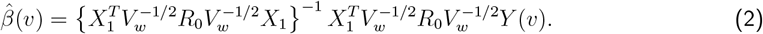

To obtain the one-step estimator for the exchangeable covariance structure, *V*_*w*_ is taken to be identity to obtain an initial estimate of *β*(*v*), then the correlation parameter, 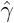, is estimated conditional on the initial estimate of *β*(*v*) and the final estimator, Zeger, 1986) (Supplementary Section S2). 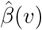, is obtained by plugging in 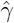 into (2) (Liang & Zeger, 1986) (Supplementary Section S2).

The robust estimator for the asymptotic covariance of 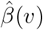, denoted by Σ_*β*_(*v, w*), is derived by a first-order Taylor expansion (Liang & Zeger, 1986). We estimate the asymptotic covariance with 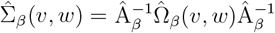, where

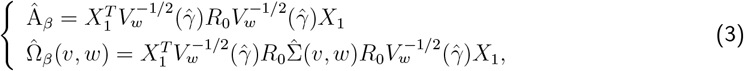

and 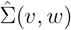 is block-diagonal with elements 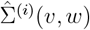, as defined in the Supplemental Equation (S2) (Long & Ervin, 2000).

Given Equations (2) and (3), the Wald test statistic can be constructed for a reference null hypothesis *H*_0_(*v*) : *β*(*v*) = *β*_0_(*v*). When the variance is known, this test statistic asymptotically follows a non-central chi-squared distribution with *m*_1_ degrees of freedom where the non-centrality parameter is 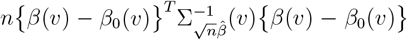, (S. Vandekar et al., 2020). When the variance of the parameter is unknown, the test statistic image is

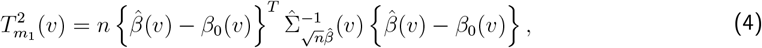

which can be used to compute the RESI (Jones et al., 2023).

### 2.3 RESI definition, estimator, and bootstrap

The RESI is based on M-estimators, such as those obtained from a GEE. Our reason for using the RESI as a standardized effect size index is that it is robust, generalizable, and easily interpretable across a wide range of models and study designs (S. Vandekar et al., 2020).

Detailed definitions and theory are given in prior work (Jones et al., 2023; Kang et al., 2023; S. Vandekar et al., 2020). To summarize, for a given voxel, *v*, the RESI is defined as the component of the noncentrality parameter of the chi-squared statistic that does not depend on the sample size

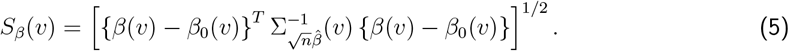

A consistent estimator of the RESI is obtained from the test statistic as (López-Blázquez, 2000; S. Vandekar et al., 2020)

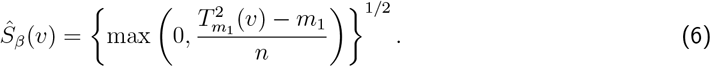

We use a nonparametric bootstrap procedure to construct confidence sets for the RESI. In general, the T-quantile nonparametric bootstrap has better small sample performance than the Normal-quantile bootstrap because it accounts for the estimation of the variance of the parameter estimator and has smaller higher-order error terms than the Normal-quantile bootstrap (Hall, 2013). However, the T-quantile bootstrap requires an estimate of the variance of the RESI estimator in each bootstrap (Hall, 2013). Because deriving the variance of (6) is challenging due to the max operator, we instead derive the asymptotic variance of 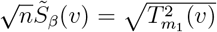 assuming the data are normally distributed and using the delta method (Supplementary Section S3),

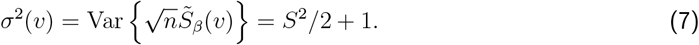

We use a plug-in estimator for the square root of this variance term to normalize the effect size estimator in each bootstrap. Although the normality assumption is likely violated, performing this normalization may improve the performance of the bootstrap if it is approximately accurate (Hall, 2013).

#### Algorithm 1

T-quantile bootstrap

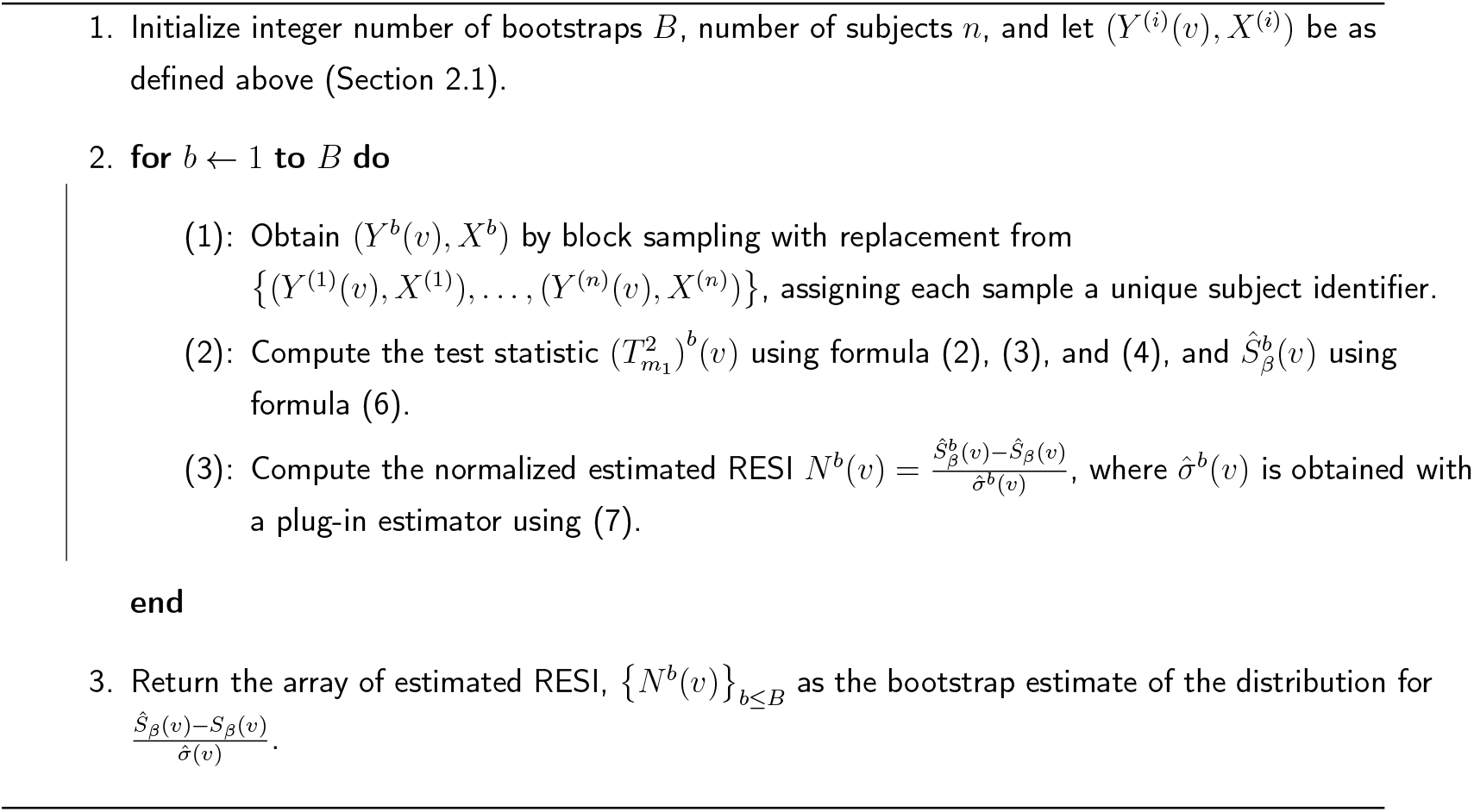

The correlation parameter *γ* in the working covariance is fixed across the bootstraps. We evaluate this T-quantile bootstrap and the Normal-quantile bootstrap, where standard deviation, 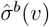, is taken to be 1, in simulations below.

### 2.4 Confidence sets

Confidence sets define a collection of voxels that form inner (or outer) spatial boundaries used to make probabilistic statements about brain regions where the true effect size is larger (or smaller) than a given non-zero effect size threshold (Bowring et al., 2021; Sommerfeld et al., 2018). For a given effect size threshold *s* and confidence level 1 − *ϵ*, the inner and outer confidence sets are defined as CS_in_(*s, ϵ*) ⊂ 𝕍 and CS_out_(*s, ϵ*) ⊂ 𝕍 that satisfy (Ren et al., 2023)

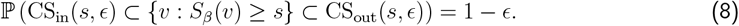

Confidence sets uniquely allow scientists to define null regions, based on effect size thresholds, as the complement of outer confidence set, CS_out_(*s, ϵ*)^*C*^, which is not possible with hypothesis testing.

Prior work by Ren et al. (2023) showed that 1 − *ϵ* level confidence sets can be derived from simultaneous confidence intervals (SCIs) in a regression context that are valid for all effect size thresholds, *s* in (8). We use their result to construct confidence sets for the RESI in longitudinal GEE using Algorithm 1 (Hall, 1992).

#### Algorithm 2

1-*ϵ* level RESI confidence set

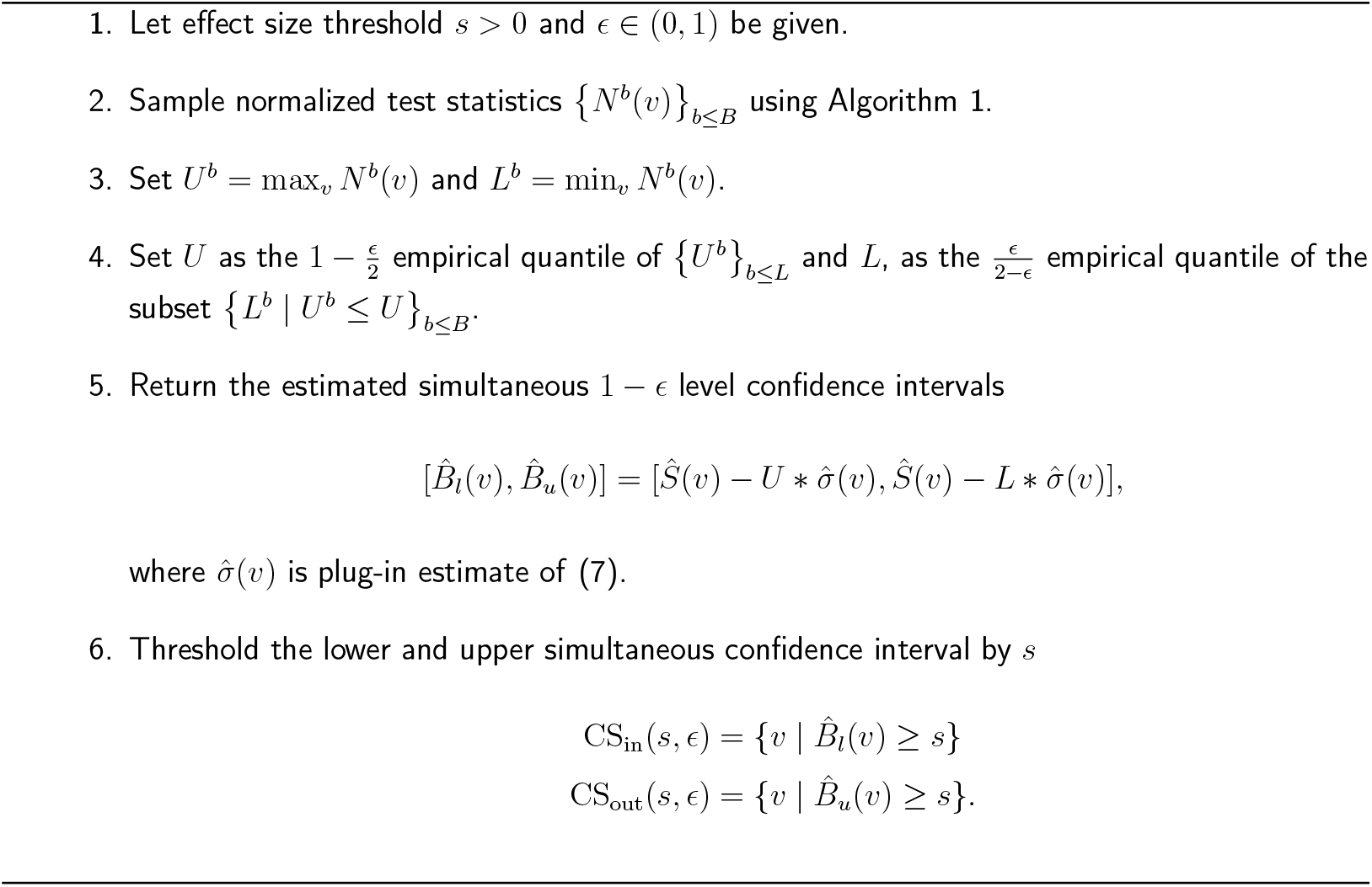

## 3 Methods

### 3.1 Simulation methods

We use simulations to evaluate the performance of the confidence set procedures under different scenarios. Because the confidence sets we use are based on simultaneous confidence intervals (Ren et al., 2023), we evaluate the bias, simultaneous and voxel-wise coverage, and width of the simultaneous confidence intervals in 500 replications. Simultaneous intervals are constructed at the 95% confidence level. This simulation is conducted on the 46th slice of the Psychiatric Genotype-Phenotype Project (PGPP; see below) data, which includes 1,845 voxels in the study mask.

For the simulations, we use a spatio-temporal covariance structure that is the Kronecker product of separate spatial and temporal covariances (Chi, 2016). Let 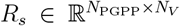 represent the spatial residuals estimated from the PGPP data using GEE (and scaled by 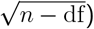, where *N*_PGPP_ is the total number of observations in the PGPP data, *n* is the number of independent participants, df is the model degrees of freedom, and *N*_*V*_ is the number of voxels in the PGPP data. The true spatial covariance matrix of the data is simulated assuming 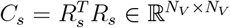. To simulate a temporal covariance between repeated observations within a participant, we let *C*_*t*_ ∈ ℝ^*N×N*^ denote the temporal correlation structure that we allow to be independent, exchangeable, or AR1, where *N* is the total number of observations across all participants in our simulation. This structure can be represented as 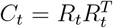, where *R*_*t*_ is derived through Cholesky decomposition. We construct the spatio-temporal error term by

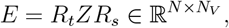

where 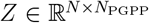 is a matrix of independent standard normal random variables. The covariance of the errors *E* is

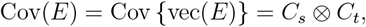

where Cov and vec are the covariance and vectorization operators.

We simulate a longitudinal regression model

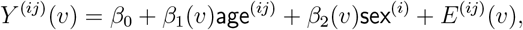

where the age^(*ij*)^ covariate (varies within-subject) is generated from the Normal distribution with a mean uniformly distributed from 20 to 70, and a variance of 1 for the *i*-th participant and the *j*-th measurement. sex^(*i*)^ (fixed within-subject) is sampled from a Bernoulli distribution with a probability of 0.5. The number of measurements per subject is randomly generated from a uniform distribution ranging between 1 and 3. The true value for 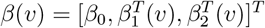 is computed from estimates of a real dataset of fractional amplitude of low-frequency fluctuations (fALFF) multiplied by scalar values (0.7 for age effect and 2 for sex effect). *β*_0_ is set to be 1. We vary the sample size *n* ∈ {25, 100, 200} across the six combinations of three true temporal covariances (independent, exchangeable, and AR1) and two working covariances (independent and exchangeable), and estimate the confidence sets using 1,000 bootstrap samples and the two Bootstrap methods mentioned in Section 2.3.

The true effect size depends on the working covariance structure as it changes the asymptotic covariance of the estimator for *β*(*v*). The asymptotic covariance 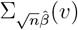 in Equation (5) is

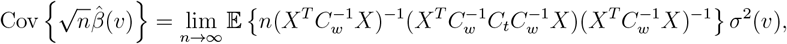

where *C*_*w*_ is the working correlation, *C*_*t*_ is the true correlation, and *σ*^2^(*v*) is the empirical variance of the residuals from the model fit in the PGPP data (Supplementary Section S4). To determine this value computationally, we use Monte Carlo integration with 1,000 replicates with 3,000 subjects to approximate this asymptotic value.

The asymptotic value of the working covariance parameter estimator is only equal to the true covariance parameter when the covariance is correctly specified. In the case where either the true or working covariance is independent, 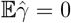. When the true covariance follows an AR1 structure, but the working covariance is assumed to be exchangeable, the asymptotic value of 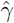 is given by (Wang, 2003):

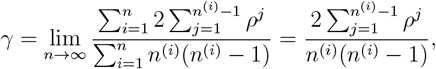

where *ρ* is the true correlation parameter in the true AR1 covariance. The proof of this result is provided in the Supplementary Section S5.

### 3.2 The ADNI dataset analysis

The Alzheimer’s Disease Neuroimaging Initiative (ADNI) is a longitudinal, multi-center, observational study. Patients were recruited across North America from 2004 through the present, the diagnostic categories include control (CN), some memory concerns (SMC), early and late mild cognitive impairment (EMCI, LMCI), and Alzheimer’s Disease (AD). Average cortical thickness within regions of the Desikan-Killiany atlas was estimated using the Freesurfer cortical parcellation pipeline. Quality control was performed using a combination of expert visual curation and automated metrics of image quality (Bethlehem et al., 2022). Participant demographic information can be found in Table 1.

**Table 1:** Participant demographic information by diagnosis in the ADNI dataset, *n* denotes the total number of participants.

| | CN<br>($n=414$) | SMC<br>($n=102$) | EMCI<br>($n=308$) | LMCI<br>($n=558$) | AD<br>($n=327$) |
| --- | --- | --- | --- | --- | --- |
| **Sex** |  |  |  |  |  |
| Female | 206<br>(49.8%) | 59<br>(57.8%) | 138<br>(44.8%) | 216<br>(38.7%) | 148<br>(45.3%) |
| Male | 208<br>(50.2%) | 43<br>(42.2%) | 170<br>(55.2%) | 342<br>(61.3%) | 179<br>(54.7%) |
| **Age (years)** |  |  |  |  |  |
| Mean | 74.8 | 72.0 | 71.1 | 74.0 | 74.9 |
| (SD) | (5.77) | (5.55) | (7.46) | (7.49) | (7.87) |
| Median | 74.1 | 71.2 | 70.8 | 74.4 | 75.4 |
| [Min, Max ] | [56.2, 89.7] | [59.7, 90.2] | [55.0, 88.7] | [54.4, 91.5] | [55.1, 91.0] |
| **Session** |  |  |  |  |  |
| 1 | 413 (26.0%) | 102 (76.1%) | 306 (27.0%) | 556 (27.9%) | 327 (43.3%) |
| 2 | 372 (23.4%) | 20 (14.9%) | 272 (24.0%) | 471 (23.6%) | 232 (30.7%) |
| 3 | 337 (21.2%) | 6 (4.5%) | 249 (22.0%) | 394 (19.8%) | 175 (23.2%) |
| 4 | 241 (15.2%) | 6 (4.5%) | 202 (17.8%) | 275 (13.8%) | 21 (2.8%) |
| 5 | 146 (9.2%) | 0 (0%) | 97 (8.6%) | 156 (7.8%) | 0 (0%) |
| 6 | 57 (3.6%) | 0 (0%) | 8 (0.7%) | 71 (3.6%) | 0 (0%) |
| 7 | 15 (0.9%) | 0 (0%) | 0 (0%) | 47 (2.4%) | 0 (0%) |
| 8 | 6 (0.4%) | 0 (0%) | 0 (0%) | 11 (0.6%) | 0 (0%) |
| 9 | 2 (0.1%) | 0 (0%) | 0 (0%) | 7 (0.4%) | 0 (0%) |
| 10 | 0 (0%) | 0 (0%) | 0 (0%) | 4 (0.2%) | 0 (0%) |

We use the proposed methods to study associations of cortical thickness with age and diagnosis (Table 1). We fit cortical thickness in each region including sex, a spline term for age with 3 degrees of freedom, a sixty-two-level factor variable for the site, and a five-level factor variable for the group with an exchangeable working correlation structure.

### 3.3 Early psychosis analysis in the PGPP

The PGPP includes data collected at the Vanderbilt Psychiatric Hospital between May 2013 and February 2018 for a prospective two-year longitudinal study, and approved by the IRB at Vanderbilt University Medical Center. The inclusion criteria for patients were a diagnosis of a non-affective psychotic disorder with a duration of psychosis less than two years. Exclusion criteria for all participants included significant head injury, major medical illnesses, pregnancy, presence of metal, claustrophobia, and current substance abuse or dependence within the past month prior to study enrollment. Participant demographic and clinical characteristics are listed in Table 2 (McHugo et al., 2022).

**Table 2:** Demographic and clinical characteristics by diagnosis for participants in the PGPP study, *n* denotes the total number of participants.

| | Early Psychosis<br>($n=59$) | Healthy Control<br>($n=67$) |
| --- | --- | --- |
| **Age** |  |  |
| Mean (SD) | 21.4 (3.93) | 21.8 (2.84) |
| Median [Min, Max] | 20.0 [16.0, 39.0] | 21.0 [17.0, 29.0] |
| **Sex** |  |  |
| Female | 13 (22.0%) | 17 (25.4%) |
| Male | 46 (78.0%) | 50 (74.6%) |
| **Race** |  |  |
| Asian | 1 (1.7%) | 1 (1.5%) |
| Black or African American | 13 (22.0%) | 10 (14.9%) |
| White | 45 (76.3%) | 53 (79.1%) |
| Other | 0 (0%) | 3 (4.5%) |
| **Time** |  |  |
| Baseline | 59 (53.6%) | 67 (56.8%) |
| Follow-up | 51 (46.4%) | 51 (43.2%) |

We used these data to perform simulation analyses and estimate effect size confidence sets for the association between diagnosis and fractional amplitude of low frequency fluctuations (fALFF) in the hippocampus. fALFF is a voxel-level frequency-domain scalar summary of brain activity during a resting-state functional MRI scan. After registration of the fMRI time series data to the MNI 2mm template, fALFF was calculated using AFNI’s 3dRSFC tool (Taylor & Saad, 2013) as the average of the power in the range 0.01–0.1 Hz relative to the entire frequency spectrum, and normalized by dividing the mean fALFF within the whole brain at each voxel (McHugo et al., 2022). Fractional ALFF is less susceptible than standard ALFF to motion, physiological artifacts, and site differences compared to standard ALFF, making it a more reliable indicator of neural activity in clinical populations (Yan et al., 2013; Zou et al., 2008). Data from nine participants were excluded for imaging data quality control or due to ineligible diagnoses at follow-up, yielding 59 patients and 67 healthy control participants. Details of image processing, data quality control, and exclusionary criteria are available in prior work (McHugo et al., 2022).

These data were used to investigate the hypothesis of progressive hippocampal excitation/inhibition (E/I) imbalance in the early stages of psychosis (McHugo et al., 2022). Our goal is to estimate confidence sets for the effect size of the group (healthy control vs. early psychosis) at all voxels in the hippocampus, with time (t0 = study entry, t2yr = two-year follow-up), sex, race, and age fit in spline included as a covariate and repeated measurements within participant modeled with an exchangeable working correlation structure.

## 4 Results

### 4.1 Simulation results

We ran simulations to assess the simultaneous coverage and mean width of the confidence intervals (averaged across all voxels) used to construct the confidence sets. The simultaneous coverage is above the nominal level and decreases to the nominal level as the sample size increases, regardless of the true or working covariance structure (Figure 1, left). As expected, confidence interval width decreases with increasing sample size, and the T-quantile bootstrap has substantially smaller confidence intervals in small samples across all scenarios (Figure 1, right). Because the T-quantile bootstrap produces narrower SCIs and has equivalent coverage it is superior to the Normal-quantile bootstrap.

**Figure 1.**
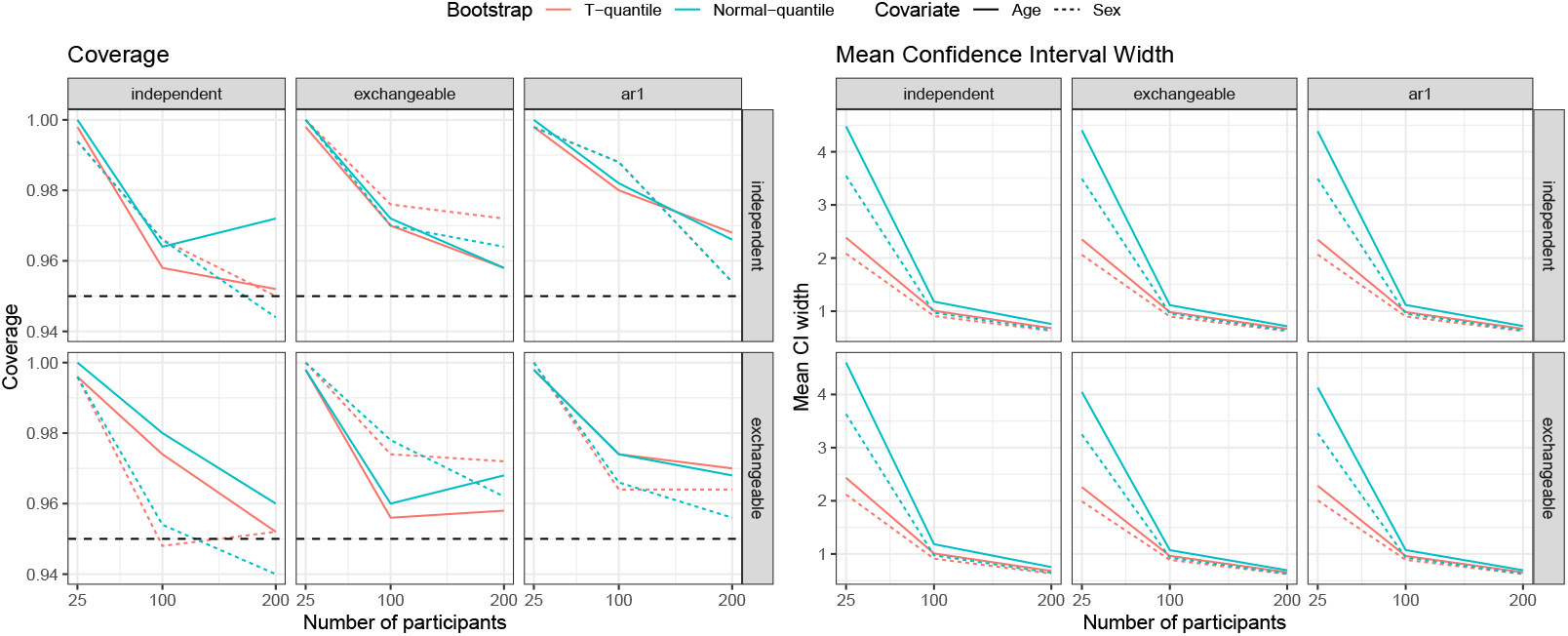
Simulation results for simultaneous coverage (left) and mean width of the simultaneous confidence intervals (right) used to construct confidence sets. The panel columns indicate the true covariance structures (independent, exchangeable, and AR1) and working covariance structures (independent and exchangeable) for different sample sizes (25, 100, and 200). Each scenario considers two covariates (age and sex) and two Bootstrap methods (T-quantile and Normal-quantile).

We also evaluated spatial patterns of bias of the effect size estimator at the voxel-level (Figure 2, top). As expected, bias decreases as the sample size increases. We evaluated voxel-wise coverage to ensure that there are no spatial patterns of false positives (Figure 2, bottom). False positives were rare and appeared randomly throughout the image plane.

**Figure 2.**
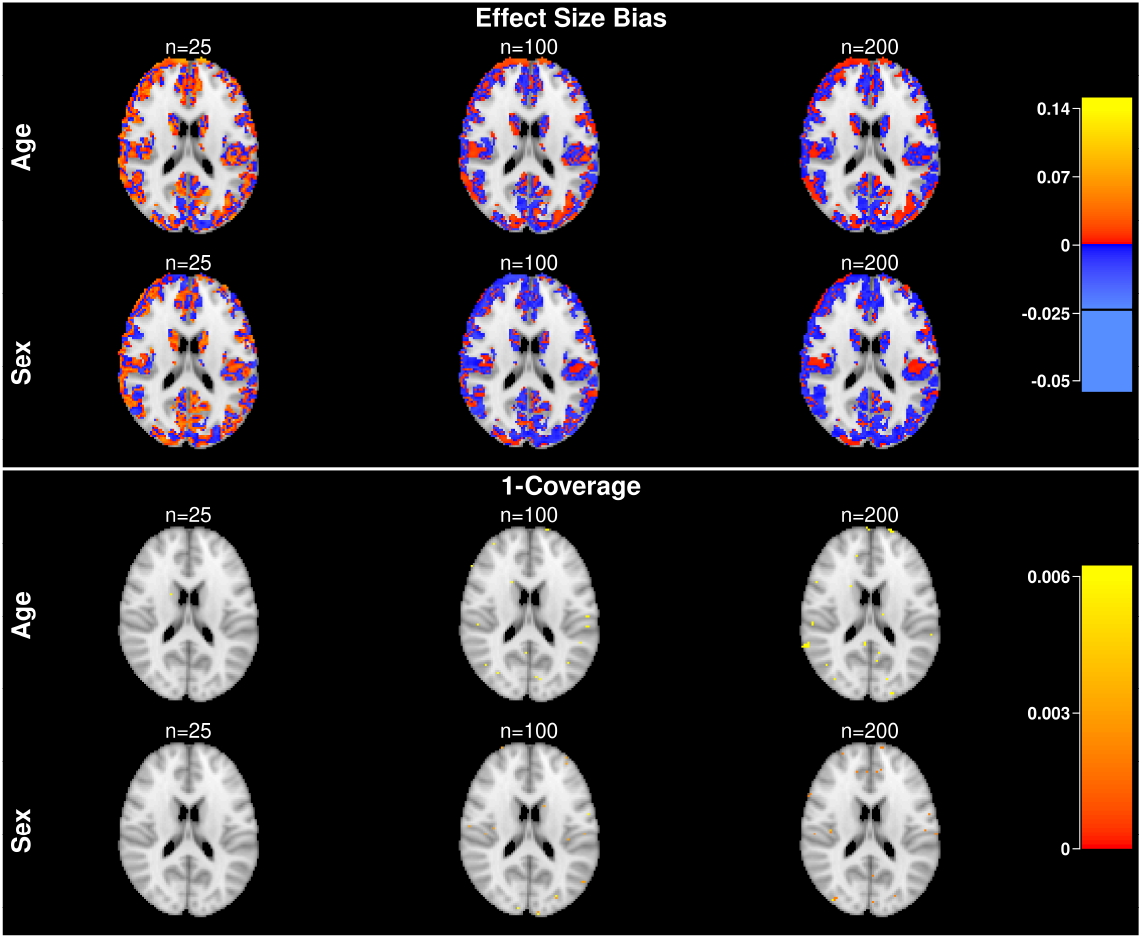
Bias of the effect size estimator (top) and 1-coverage (bottom) when the true covariance structure is AR1 and working structure is exchangeable. The range of the color bar is taken from the 90% percentile of the maximum and minimum values across six scenarios with three different sample sizes and two covariates.

### 4.2 Cortical thickness in Azheimer’s Disease

We use the proposed methods to construct confidence sets of the RESI for associations of age and the five diagnostic groups in the longitudinal ADNI dataset within the Desikan-Killany atlas. We visualize regions in the “null set”, {CS_out_(*s, ϵ*)}^*C*^, where the effect size is likely to be smaller than a given threshold, *s*, in blue/cyan and regions in the target set, CS_in_(*s, ϵ*), in red/yellow. The lower simultaneous confidence limit is used to construct the target set and the upper confidence limit is used to construct the null set (Figure 3).

**Figure 3.**
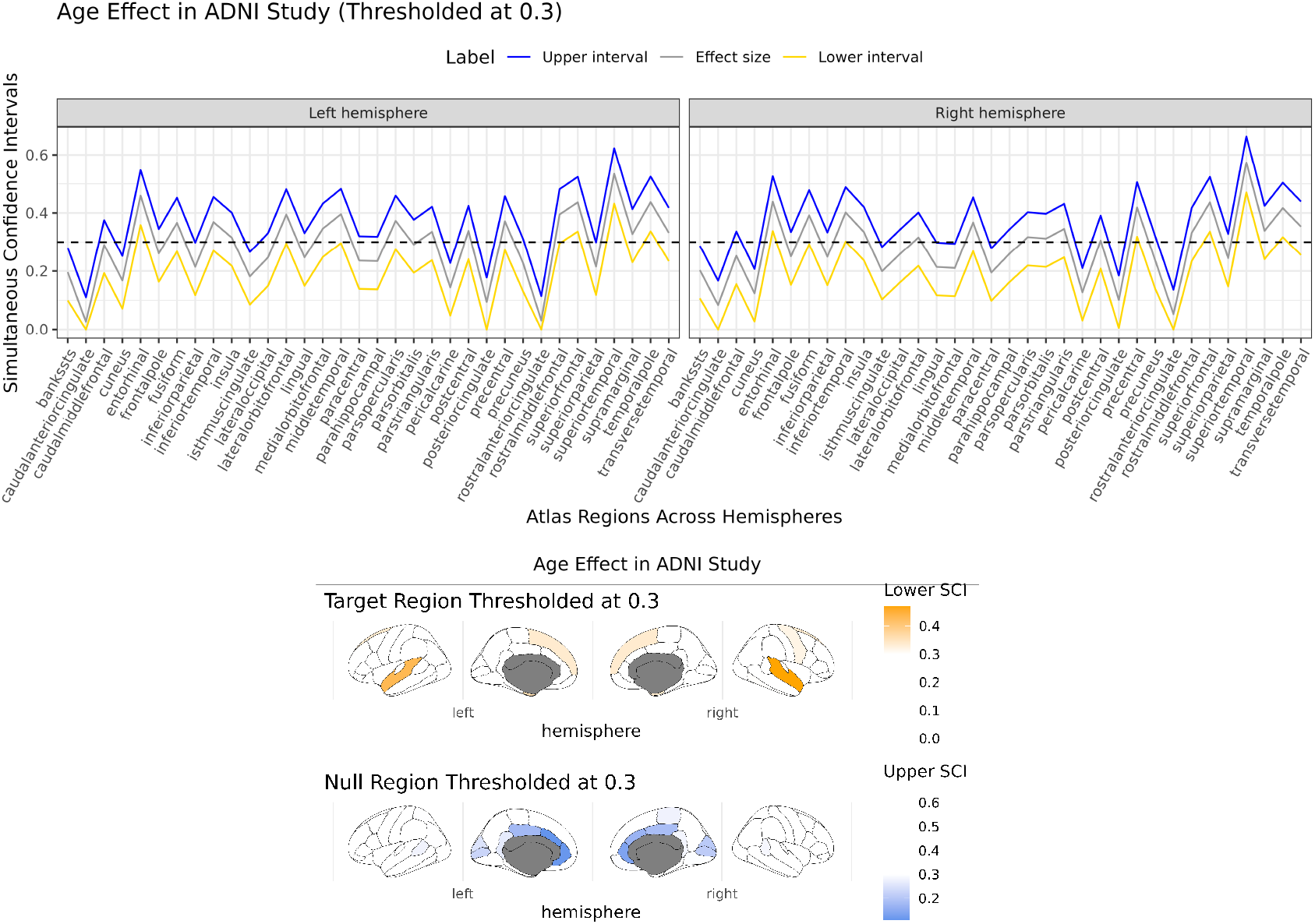
Simultaneous confidence intervals (top) and confidence sets (bottom) for the associations of cortical thickness with age RESIs across all atlas regions in the left and right hemispheres. The lower confidence limit (yellow) is used to construct an inner confidence set using the threshold *s* = 0.3 (dashed line), which identifies regions of the brain where the true effect size is likely to be larger than that value. The upper confidence limit is used to construct the null set using the same threshold, identifying regions where the effect size is likely to be smaller than that value (Section 2.4).

We calculated the RESI for the age and diagnosis effects corresponding to each atlas region along with the simultaneous confidence intervals. We chose *s* = 0.3 as the threshold corresponding to a “medium” effect size using Cohen’s recommendations (Cohen, 1988) and used confidence sets to identify regions whose association is stronger than 0.3 with 95% confidence (in the Frequentist sense) by thresholding the lower limit, and regions with effect sizes larger than this threshold using the upper confidence limit. Regions above this limit could serve as biomarkers of normative aging or to distinguish the patient groups. Because the confidence sets are simultaneous we can apply them at other thresholds and still maintain the same error rate (Ren et al., 2023).

For age, the superior temporal lobe, superior frontal lobe, and entorhinal cortex have effect sizes larger than 0.3 with 95% confidence indicating at least medium strength associations of cortical thickness with natural aging within these regions (Figure 3, bottom, “Target Region”). Regions in the anterior cingulate cortex, occipital lobe, and motor cortex were identified as being in the null set (using *s* = 0.3 threshold), identifying where the effect size is confidently less than a “medium” effect size, highlighting regions in the occipital lobe and anterior cingulate cortex (ACC). We have 95% confidence over the whole brain that all regions inside the null set have a true RESI of less than 0.3. Notably, most of the simultaneous confidence intervals do not contain zero, indicating that we would reject the null in most regions. The confidence set approach allows us to identify important regions based on their effect size instead of the hypothesis test.

For diagnosis, we identify temporal, parietal, and motor regions with effect sizes larger than *s* = 0.3, with strongest effect sizes middle temporal, superior temporal, and entorhinal cortex (Figure 4, bottom, “Target Region”). These regions of strongest effect sizes align with the wide body of research implicating the temporal lobe in memory decline and AD (Chan et al., 2001; Visser et al., 1999). We defined a null region using the upper interval of 0.3 capturing regions in the ACC and inferior frontal lobe. Although highlighting only a few regions, this set identifies regions that are very likely to have a smaller than “medium” effect size. Taken together, these results suggest the largest diagnostic effects are isolated in the temporal and parietal lobes. As with the age effect, most simultaneous intervals do not include zero, indicating that adjusted hypothesis tests would identify most regions as significant. The confidence set approach allows investigators to identify regions as important based on their effect size instead.

**Figure 4.**
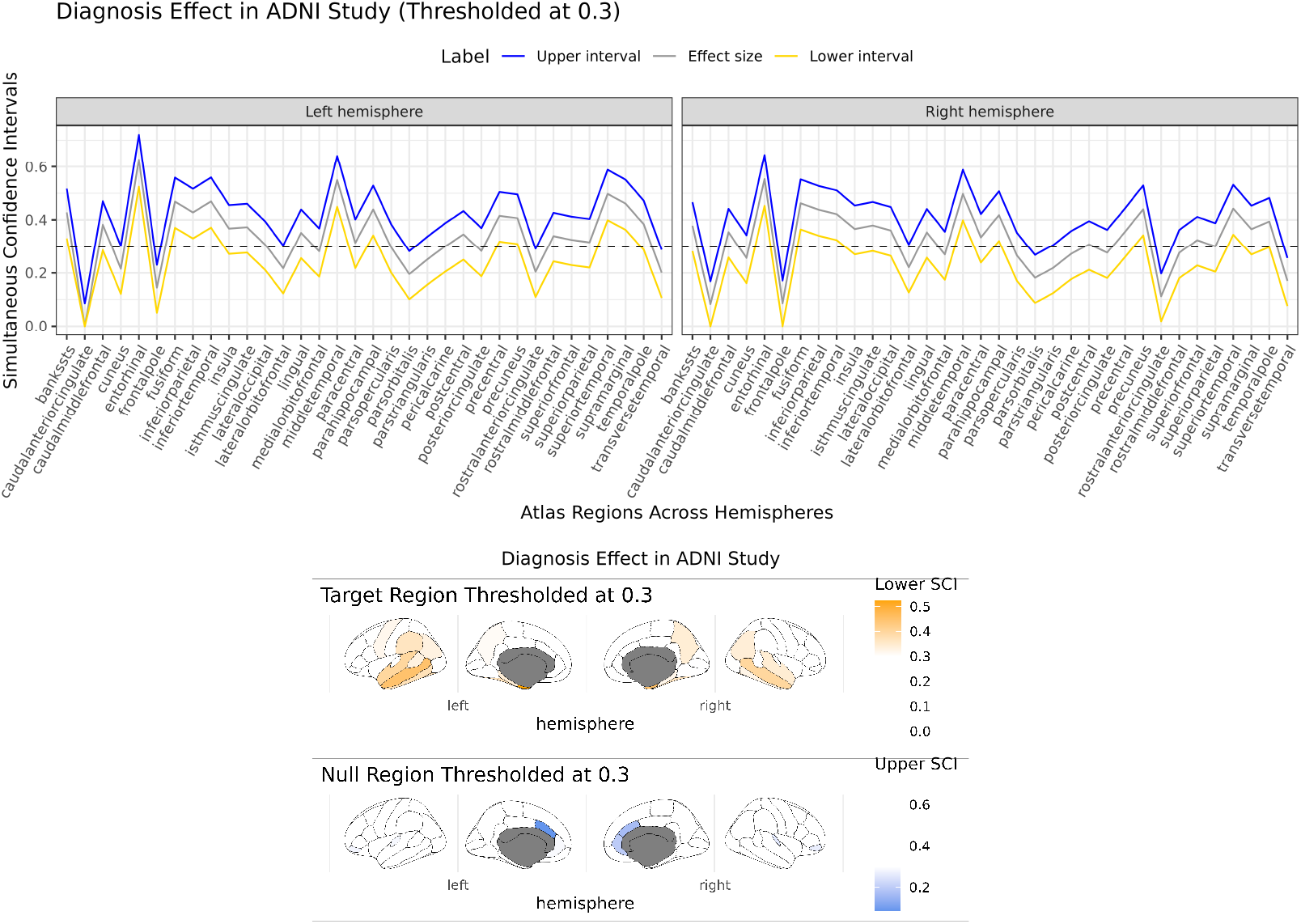
Simultaneous confidence intervals (top) and confidence sets (bottom) for the associations of cortical thickness with diagnosis RESIs across all atlas regions in the left and right hemispheres. The lower confidence limit (yellow) is used to construct an inner confidence set using the threshold *s* = 0.3 (dashed line), which identifies regions of the brain where the true effect size is likely to be larger than that value. The upper confidence limit is used to construct the null set using the same threshold, identifying regions where the effect size is likely to be smaller than that value (Section 2.4).

### 4.3 Differences in fALFF in early psychosis

We investigated the association between early psychosis and fALFF in the hippocampus using the proposed methods. The simultaneous confidence intervals for the association of early psychosis with fALFF show no voxels have a lower confidence limit above zero (Figure 5, top) suggesting that there are limited regions where we can confidently conclude an effect of early psychosis on hippocampal E/I imbalance. Thresholding the upper simultaneous confidence limit below 0.4 identifies regions where the effect size is likely to be smaller than a “large” effect size (Cohen, 1988; S. N. Vandekar et al., 2019) (Figure 5, bottom). One strength of this approach is to systematically exclude these regions from further analysis, for example, if investigators are searching for large effect sizes for the sake of establishing biomarkers.

**Figure 5.**
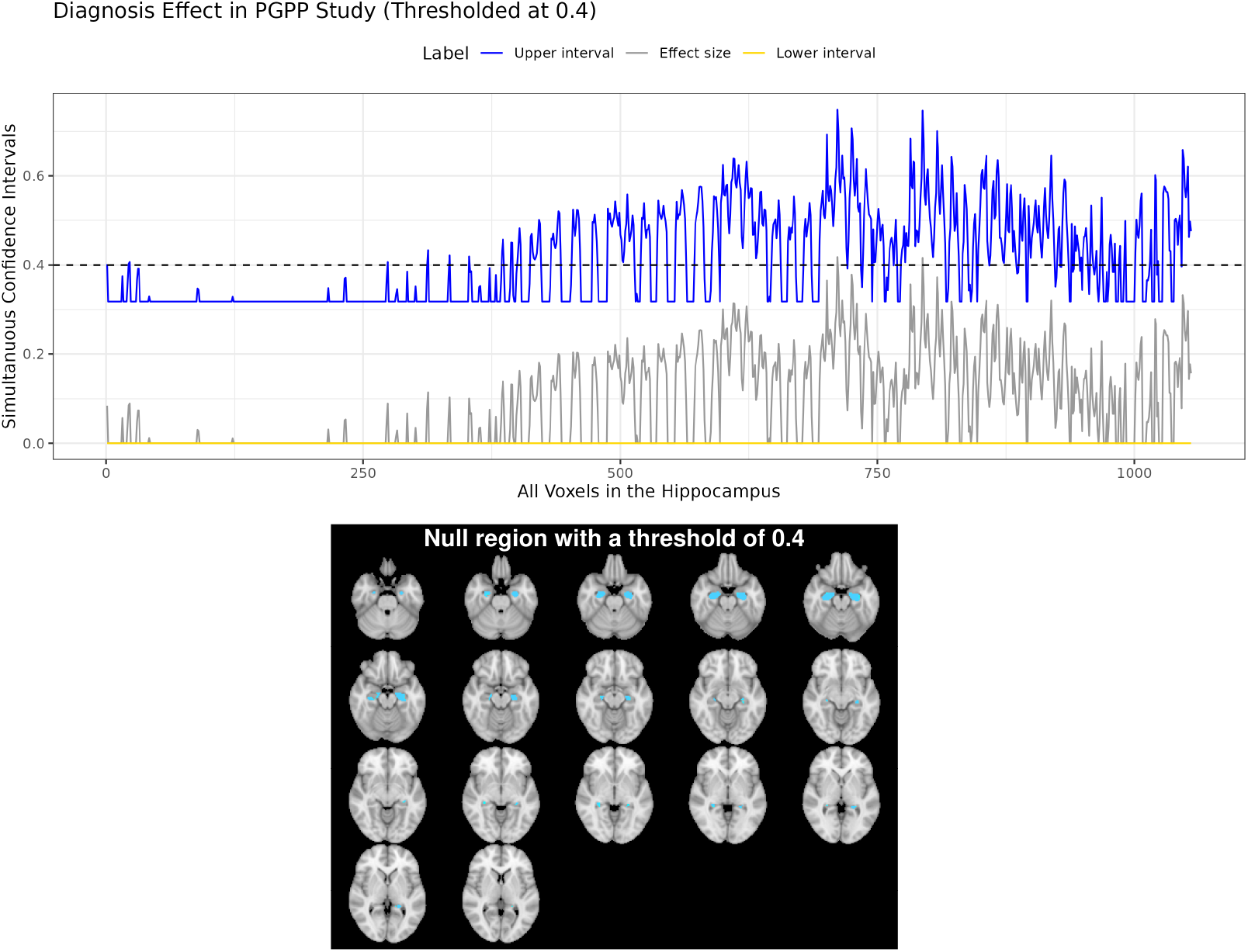
Simultaneous confidence intervals (top) and null sets (bottom) for the associations of fALFF with diagnosis RESIs across all voxels in the hippocampus. The upper confidence limit (blue) is used to construct the null set using a threshold of *s* = 0.4 (dashed line), identifying regions where the true effect size is likely to be below a “large” effect size (Section 2.4).

By identifying “null” regions with 95% confidence, we can avoid unnecessary examination of these areas, improving efficiency in future studies. As above, the use of simultaneous intervals allows us to visualize the confidence sets at multiple thresholds without increasing the error rate.

## 5 Discussion

We developed an approach to construct confidence sets for arbitrary effect sizes in cross-sectional and longitudinal analyses and used the methods to study diagnostic associations in Alzheimer’s disease with cortical thickness and associations of psychosis-spectrum diagnosis with resting state activity in the hippocampus. Our methods allowed us to perform analyses for multiparameter statistics, such as nonlinear age and multi-group diagnoses in longitudinal models – these analyses were not possible with existing methods for confidence sets (Bowring et al., 2019, 2021; Sommerfeld et al., 2018). In contrast to hypothesis testing approaches for neuroimaging data, confidence sets can be used to identify regions likely to be in a null set, defined by having a true effect size below a given value. Constructing confidence sets using simultaneous confidence intervals allows the user to construct confidence sets across all possible effect size thresholds while still maintaining 95% coverage.

A critical limitation of the method is that the width of the intervals can be wide for small studies. Using the simultaneous intervals is similar to controlling the voxel-wise family-wise error rate using hypothesis testing, which can lead to conservative results. For example, in the PGPP study, which included 126 participants, the intervals were too large to identify regions in the target set. This limitation is noted by Bowring et al. (2021) when constructing confidence sets for Cohen’s *d*. For this reason, the confidence set approach may be better suited for larger neuroimaging studies where effect size estimation is the primary objective. Potentially, meta-analytic methods can be used to aggregate effect estimates across studies to build tighter confidence sets. It may also be possible to combine our work with theoretically optimal methods for constructing simultaneous confidence intervals (Gao et al., 2021) to construct tighter confidence sets. In addition, developing methods of confidence sets for a single, fixed threshold, or that control the equivalent of the false discovery rate may lead to tighter confidence sets (Ren et al., 2023; Sommerfeld et al., 2018).

In conclusion, the methods developed here generalize confidence sets to arbitrary effect sizes and longitudinal data, significantly expanding the research questions that can be studied using confidence sets. With the code available in the pbj R package, these methods can improve effect size reporting in neuroimaging.

## Data and Code Availability

The ADNI Data used in this article were obtained from the Alzheimer’s Disease Neuroimaging Initiative (ADNI) database (adni.loni.usc.edu). The PGPP data are not publicly available due to ethical restrictions. The methods presented in this paper are built upon functions provided by the pbj R package (https://github.com/statimagcoll/pbj), and the codes for analyses can be found here (https://github.com/statimagcoll/Confidence_Sets.git).

## Author Contributions

Xinyu Zhang: Conceptualization, Methodology, Software, Formal analysis, Writing—original draft, Writing—Review & Editing Kenneth Liao: Conceptualization, Methodology, Software, Writing—Review & Editing Jakob Seidlitz: Data Curation, Investigation, Writing—Review & Editing Maureen McHugo: Data Curation, Investigation, Writing—Review & Editing Suzanne N. Avery: Data Curation, Investigation, Writing—Review & Editing Anna Huang: Data Curation, Investigation, Writing—Review & Editing Aaron Alexander-Bloch: Investigation, Writing—Review & Editing Neil Woodward: Investigation, Writing—Review & Editing, Funding acquisition Stephan Heckers: Investigation, Writing—Review & Editing, Funding acquisition Simon Vandekar: Conceptualization, Methodology, Software, Writing—Review & Editing, Supervision, Validation, Funding acquisition

## Acknowledgements

The ADNI was launched in 2003 as a public-private partnership, led by Principal Investigator Michael W. Weiner, MD. The primary goal of ADNI has been to test whether serial magnetic resonance imaging (MRI), positron emission tomography (PET), other biological markers, and clinical and neuropsychological assessment can be combined to measure the progression of mild cognitive impairment (MCI) and early Alzheimer’s disease (AD). For up-to-date information, see www.adni-info.org.

## Supplementary Material

Supplementary material is available.

