## Supplementary materials for "Semiparametric Confidence Sets for Arbitrary Effect Sizes in Longitudinal Neuroimaging"

February 4, 2025

#### S1 Residual Forming Matrix

We define the residual forming matrices as,

$$\begin{aligned} R_0 &= I - V_w^{-1/2} X_0 \left( X_0^T V_w^{-1} X_0 \right)^{-1} X_0^T V_w^{-1/2}, \\ R &= I - V_w^{-1/2} X \left( X^T V_w^{-1} X \right)^{-1} X^T V_w^{-1/2}, \end{aligned} \tag{S1}$$

which project vectors onto the orthogonal complement of the column space of the design matrices $X_0$ and $X$, respectively.

Subject-level temporal covariances are estimated by Long and Ervin, 2000

$$\hat{\Sigma}^{(i)}(v, w) = R_{i,i}^{-1} \left\{ R V_w^{-1/2} Y(v) \right\}^{(i)} \left\{ R V_w^{-1/2} Y(w) \right\}^{(i)T} R_{i,i}^{-1}. \tag{S2}$$

### S2 Working Covariance Structure

Let $V_w^{(i)}(\gamma) = \big(A^{(i)}\big)^{1/2} C_w^{(i)}(\gamma) \big(A^{(i)}\big)^{1/2} \in \mathbb{R}^{n^{(i)} \times n^{(i)}}$ represent the working covariance matrix for the $i$-th subject. Here, $\big(A^{(i)}\big)^{1/2} = \mathrm{diag}\{\sigma^{(i)}\}$ contains the standard deviations of $Y^{(i)}$, and $C_w^{(i)}(\gamma)$ denotes the working correlation matrix of the $n^{(i)}$ repeated measures, parameterized by $\gamma$. Computing $\big(V_w^{(i)}(\gamma)\big)^{-1}$ requires inverting $C_w^{(i)}(\gamma)$, which can be computationally challenging.

#### S2.1 The Exchangeable Structure

The exchangeable correlation structure for the $i$-th subject is defined as

$$C_w^{(i)}(\gamma) = \begin{bmatrix} 1 & \gamma & \cdots & \gamma \\ \gamma & 1 & \cdots & \gamma \\ \vdots & & \ddots & \vdots \\ \gamma & \cdots & \gamma & 1 \end{bmatrix},$$

and we assume all subjects share the same correlation parameter $\gamma$. Following Example 3 in Liang and Zeger, 1986, the parameter $\gamma$ can be estimated using the Pearson residuals by

$$\begin{aligned} \hat{\gamma} &= \left\{ \sum_{i=1}^{n} \frac{n^{(i)}(n^{(i)}-1)}{2} - m_1 \right\}^{-1} \sum_{i=1}^{n} \sum_{j<k} \hat{e}^{(ij)} \hat{e}^{(ik)} \\ &= \left\{ \sum_{i=1}^{n} \frac{n^{(i)}(n^{(i)}-1)}{2} - m_1 \right\}^{-1} \sum_{i=1}^{n} \frac{\left( \sum_{j=1}^{n^{(i)}} \hat{e}^{(ij)} \right)^2 - \sum_{j=1}^{n^{(i)}} \left( \hat{e}^{(ij)} \right)^2}{2}, \end{aligned}$$

where $m_1$ is the number of covariates in $X_1$, $n$ is the number of subjects, and $n^{(i)}$ is the number of repeated measures for subject $i$.

The Pearson residuals $\hat{e}^{(ij)}$ for the $j$-th measurement of the $i$-th subject are defined as

$$\hat{e}^{(ij)} = \frac{Y^{(ij)} - \hat{Y}^{(ij)}}{\hat{\sigma}^{(i)}},$$

where $\hat{Y}^{(ij)}$ is the fitted value for $Y^{(ij)}$, and $\hat{\sigma}^{(i)}$ is the estimated standard deviation for subject $i$, computed as

$$\hat{\sigma}^{(i)} = \left\{ \frac{\sum_{j=1}^{n^{(i)}} \left( Y^{(ij)} - \hat{Y}^{(ij)} \right)^2}{n^{(i)} - m_1} \right\}^{1/2}.$$

Then the inverse of $C_w^{(i)}(\hat{\gamma})$ can be written as (Qu et al., 2000)

$$\left\{C_w^{(i)}(\hat{\gamma})\right\}^{-1} = \frac{1}{1-\hat{\gamma}} \cdot I_{n^{(i)}} - \frac{\hat{\gamma}}{(1-\hat{\gamma})(1+(n^{(i)}-1)\hat{\gamma})} \cdot J_{n^{(i)}} = aI_{n^{(i)}} + bJ_{n^{(i)}},$$

where $I_{n^{(i)}}$ is a $n^{(i)} \times n^{(i)}$ identity matrix, $J_{n^{(i)}}$ is a matrix of one with size $n^{(i)}$,

$$a = \frac{1}{1-\hat{\gamma}},$$

$$b = -\frac{a\hat{\gamma}}{1+(n^{(i)}-1)\hat{\gamma}}.$$

To simplify notation, we omit the explicit dependence of $C_w$ on $\hat{\gamma}$, $a$, and $b$ in the equations below. To efficiently compute $\left(C_w^{(i)}\right)^{-1/2}$, we derive its explicit expression. First, consider the eigendecomposition of $\left(C_w^{(i)}\right)^{-1}$, given by

$$\left(C_w^{(i)}\right)^{-1} = U^{(i)} D^{(i)} U^{(i)T},$$

where $D^{(i)}$ is the diagonal matrix of eigenvalues, and $U^{(i)}$ contains the corresponding eigenvectors. The eigenvalues in $D^{(i)}$ are $a$ with multiplicity $n^{(i)}-1$, and $a + n^{(i)}b$ with multiplicity 1, then $D^{(i)} = \operatorname{diag}\left\{\underbrace{a, a, \ldots, a}_{n^{(i)}-1}, a + n^{(i)}b\right\}$. The eigenvectors in $U^{(i)}$ are derived via the Gram-Schmidt orthogonalization process to ensure they form an orthonormal basis,

$$v_1 = \frac{1}{\sqrt{2}}\begin{bmatrix}1\\-1\\0\\0\\\vdots\\0\\0\end{bmatrix}, v_2 = \frac{1}{\sqrt{6}}\begin{bmatrix}1\\1\\-2\\0\\\vdots\\0\\0\end{bmatrix}, \cdots, v_{n^{(i)}-1} = \frac{1}{\sqrt{n^{(i)}(n^{(i)}-1)}}\begin{bmatrix}1\\1\\1\\1\\\vdots\\-(n^{(i)}-1)\\0\end{bmatrix}, v_{n^{(i)}} = \frac{1}{\sqrt{n^{(i)}}}\begin{bmatrix}1\\1\\1\\1\\\vdots\\1\\1\end{bmatrix}.$$

The inverse square root of $C^{(i)}$, denoted as $\left(C^{(i)}\right)^{-1/2}$, can then be efficiently computed as:

$$\left(C^{(i)}\right)^{-1/2} = U^{(i)}\left(D^{(i)}\right)^{1/2} U^{(i)T}.$$

Here, $\left(D^{(i)}\right)^{1/2}$ is the diagonal matrix whose entries are the square roots of the reciprocals of the eigenvalues in $D^{(i)}$.

### S3 Asymptotic Variance of RESI Estimator

The estimator of the effect size used to derive the asymptotic variance is given by

$$\tilde{S}_\beta^2(v) = \frac{T_{m_1}^2(v)}{n}.$$

By the relationship between the non-central chi-squared distribution and the non-central $F$-distribution, as $n \to \infty$, we have

$$\frac{T_{m_1}^2(v)}{m_1} \sim \frac{\chi_{m_1}^2(\lambda)}{m_1} \propto F_{m_1,n-m}(nS_\beta^2),$$

where $nS_\beta^2$ is the non-centrality parameter (Kang et al., 2023). This implies that $T_{m_1}^2(v) \propto m_1 F_{m_1,n-m}$. Plugging it into the effect size estimator, we obtain

$$\tilde{S}_\beta^2(v) = \frac{m_1 F_{m_1,n-m}}{n}.$$

By the variance of the non-central F distribution, the asymptotic variance of $\tilde{S}_\beta^2(v)$ can be estimated as

$$\begin{aligned}
\text{Var}\left\{\tilde{S}_\beta^2(v)\right\} &= \text{Var}\left\{\frac{m_1 F_{m_1,n-m}}{n}\right\} \\
&= \left(\frac{m_1}{n}\right)^2 \text{Var}\left\{F_{m_1,n-m}\right\} \\
&= \left(\frac{m_1}{n}\right)^2 2\frac{(m_1+\lambda)^2 + (m_1+2\lambda)(n-m-2)}{(n-m-2)^2(n-m-4)} \left(\frac{n-m}{m_1}\right)^2 \\
&= 2\frac{(m_1+\lambda)^2 + (m_1+2\lambda)(n-m-2)}{(n-m-2)^2(n-m-4)} \times \frac{(n-m)^2}{n^2} \\
&\to \frac{2(S^4+2S^2)}{n} \quad (\text{as } n \to \infty).
\end{aligned}$$

Assuming data is normally distributed, by delta method, we can compute the asymptotic variance of $\tilde{S}_\beta(v)$ as

$$\text{Var}\left\{\tilde{S}_\beta(v)\right\} = \frac{S^2/2+1}{n}.$$

### S4 Asymptotic Covariance Computation

According to the law of total variance,

$$\begin{aligned}
\mathrm{Cov}\left\{\sqrt{n}\hat{\beta}(v)\right\} &= \mathbb{E}\left\{\mathrm{Cov}\left(\sqrt{n}\hat{\beta}(v) \mid X\right)\right\} + \mathsf{Var}\left\{\mathbb{E}\left(\sqrt{n}\hat{\beta}(v) \mid X\right)\right\} \\
&= \mathbb{E}\left\{\mathrm{Cov}\left(\sqrt{n}\hat{\beta}(v) \mid X\right)\right\} + \mathsf{Var}\left\{\sqrt{n}\beta(v)\right\} \\
&= \mathbb{E}\left\{\mathrm{Cov}\left(\sqrt{n}\hat{\beta}(v) \mid X\right)\right\} \\
&= n\mathbb{E}\left\{(X^T V_w^{-1} X)^{-1}(X^T V_w^{-1}\Sigma(v) V_w^{-1} X)(X^T V_w^{-1} X)^{-1}\right\} \\
&= n\mathbb{E}\left\{(X^T C_{work}^{-1} X)^{-1}(X^T C_{work}^{-1} C_{true} C_{work}^{-1} X)(X^T C_{work}^{-1} X)^{-1}\right\}\sigma^2(v),
\end{aligned}$$

### S5 Proof for Correlation Estimation

Given that we denote $\rho$ as the correlation parameter in the true covariance structure and $\gamma_{jk}$ as the working correlation between the $j$-th and $k$-th repeated measurements within each subject, we can express the general moment-based estimator for exchangeable working covariance as (Wang, 2003):

$$\sum_{i=1}^{n}\sum_{j\neq k} e^{(ij)}e^{(ik)} = \sum_{i=1}^{n}\sum_{j<k}\gamma_{jk}\left\{\left(e^{(ij)}\right)^2 + \left(e^{(ik)}\right)^2\right\},$$

where $i$ indexes the subjects from 1 to $n$, $j$ and $k$ denote repeated measurements, and $e$ represents standardized residuals. In this scenario, $\gamma_{jk} = \gamma$, which remains constant across all pairs of residuals. The equation then simplifies to:

$$\hat{\gamma} = \lim_{n\to\infty}\frac{\sum_{i=1}^{n}\sum_{j\neq k} e^{(ij)}e^{(ik)}}{\sum_{i=1}^{n}(n^{(i)}-1)\sum_{j=1}^{n^{(i)}}\left(e^{(ij)}\right)^2},$$

where $n^{(i)}$ is the number of repeated measurements for subject $i$.

Assuming that the true temporal correlation structure is independent (i.e., there is no correlation between repeated measurements within subjects in the true structure), the expected value of each pairwise product $\mathbb{E}\left\{e^{(ij)}e^{(ik)}\right\}$ for $j \neq k$ approaches zero as the sample size increases. Consequently, the numerator in the expression for $\hat{\gamma}$ converges to zero.

Assuming the true temporal correlation structure follows an AR1 process, the covariance between two observations $Y_{ij}$ and $Y_{ik}$ at time points $j$ and $k$ for subject $i$ is given by $\mathrm{Cov}\left\{Y^{(ij)}, Y^{(ik)}\right\} = e^{(ij)}e^{(ik)} = \rho^{|j-k|}$. The standardized residuals $e^{(ij)}$ are assumed to have unit variance, i.e., $\mathsf{Var}\left\{Y^{(ij)}\right\} = \left(e^{(ij)}\right)^2 = 1$.

To estimate the working correlation parameter $\gamma$ under the exchangeable assumption, we use the moment-based estimator:

$$
\begin{aligned}
\hat{\gamma} &= \lim_{n\to\infty} \frac{\sum_{i=1}^{n}\sum_{j\neq k} e^{(ij)}e^{(ik)}}{\sum_{i=1}^{n}(n^{(i)}-1)\sum_{j=1}^{n^{(i)}}\left(e^{(ij)}\right)^2} \\
&= \lim_{n\to\infty} \frac{\sum_{i=1}^{n}\sum_{j\neq k} e^{(ij)}e^{(ik)}}{\sum_{i=1}^{n} n^{(i)}(n^{(i)}-1)} \\
&= \lim_{n\to\infty} \frac{\sum_{i=1}^{n} 2\sum_{j<k} \rho^{|j-k|}}{\sum_{i=1}^{n} n^{(i)}(n^{(i)}-1)} \\
&= \lim_{n\to\infty} \frac{\sum_{i=1}^{n} 2\sum_{j=1}^{n^{(i)}-1} \rho^{j}}{\sum_{i=1}^{n} n^{(i)}(n^{(i)}-1)} \\
&= \frac{2\sum_{j=1}^{n^{(i)}-1} \rho^{j}}{n^{(i)}(n^{(i)}-1)}
\end{aligned}
$$

indicating that the estimator $\hat{\gamma}$ incorporates the decay of correlation with increasing lag, as expected under the AR1 structure. The estimator from Liang and Zeger, 1986, $\frac{\sum^{(i)}\sum_{j\neq k}\epsilon_{ij}\epsilon_{ik}}{\sum^{(i)} n^{(i)}(n^{(i)}-1)-m_1}$, has the same limit.

Another estimator is used for an AR1 working correlation, defined as

$$
\hat{\gamma} = \lim_{n\to\infty} \frac{\sum_{i=1}^{n}\sum_{j=2}^{n^{(i)}} e^{(ij)}e^{(i,j-1)}}{\sum_{i=1}^{n}\left[\sum_{j=2}^{n^{(i)}-1}\left(e^{(ij)}\right)^2 + \frac{1}{2}\left\{\left(e^{(i1)}\right)^2 + \left(e^{(in^{(i)})}\right)^2\right\}\right]},
$$

yields an expectation of zero for the numerator under an independent true covariance structure:

$$
\mathbb{E}\left\{\sum_{i=1}^{n}\sum_{j=2}^{n^{(i)}} e^{(ij)}e^{(i,j-1)}\right\} = 0,
$$

since $\mathbb{E}\left\{e^{(ij)}e^{(i,j-1)}\right\} = 0$ for all $i$ under independence. The denominator approaches $\sum_{i=1}^{n}(n^{(i)}-1)$ as $\mathbb{E}\left\{(e^{(ij)})^2\right\} = 1$ for standardized residuals. Thus, as $n\to\infty$:

$$
\hat{\gamma} \approx \frac{0}{\sum_{i=1}^{n}(n^{(i)}-1)} = 0.
$$

### S6 Simulation SCI Plots

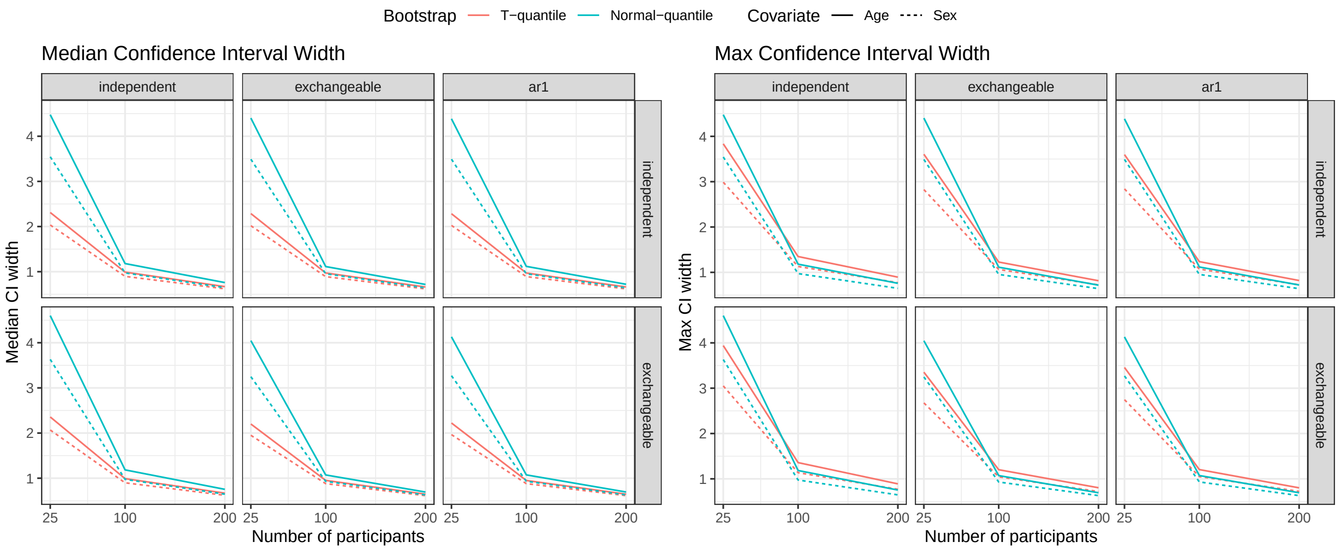


Figure S1: Median and max width of confidence sets.
